# Accurate estimation of molecular counts in droplet-based single-cell RNA-seq experiments

**DOI:** 10.1101/171496

**Authors:** Viktor Petukhov, Jimin Guo, Ninib Baryawno, Nicolas Severe, David Scadden, Maria G. Samsonova, Peter V. Kharchenko

## Abstract

Single-cell RNA-seq protocols provide powerful means for examining the gamut of cell types and transcriptional states that comprise complex biological tissues. Recently-developed approaches based on droplet microfluidics, such as inDrop or Drop-seq, use massively multiplexed barcoding to enable simultaneous measurements of transcriptomes for thousands of individual cells. The increasing complexity of such data also creates challenges for subsequent computational processing and troubleshooting of these experiments, with few software options currently available. Here we describe a flexible pipeline for processing droplet-based transcriptome data that implements barcode corrections, classification of cell quality, and diagnostic information about the droplet libraries. We introduce advanced methods for correcting composition bias and sequencing errors affecting cellular and molecular barcodes to provide more accurate estimates of molecular counts in individual cells.

## Background

RNA-seq protocols have been optimized to enable large-scale transcriptional profiling of individual cells. Such single-cell measurements require both improved molecular techniques, as well as effective ways to isolate and process a large number of cells in parallel. While single-cell RNA-seq (scRNA-seq) remains a challenging technique, several solutions are being increasingly applied, most notably techniques based on droplet microfluidics such as inDrop [1], Drop-seq [2] and 10X Chromium platform. In these approaches, cells are encapsulated in water-based droplets together with barcoded beads and necessary reagents within an oil-based flow. This allows the RNA material extracted from each cell to be contained within the droplet and tagged by a unique cellular barcode (CB) carried on the bead.

InDrop and similar approaches pool material from different cells to prepare the library, and rely on computational analysis to recognize the reads originating from the same cell based on the CB contained in the read sequence. The reads also carry a random barcode - a unique molecular identifier (UMI) [3, 4] - that can be used to discount the redundant contribution of reads originating from the same cdNa molecule as a result of library amplification. As such, the primary aim of the data processing pipeline, including the one presented here, is to provide accurate estimates of the number of molecules that have been observed for each gene in each measured cell - a molecular count matrix. Accurate estimation of such matrix is crucial, as it commonly provides the starting point for all downstream analysis, such as cell clustering or tracing of cell trajectories.

Several factors complicate the estimation of this molecular count matrix, well beyond simple parsing of the read sequences. First, the procedure must separate reads originating from droplets containing real cells from contribution of empty droplets which can amplify extracellular background transcripts and significantly outnumber the real cells. Some of the droplets may contain damaged or fragmented cells, which complicates such separation. The procedure must also address problems stemming from sequencing errors, particularly errors within the CB or UMIs which result in misclassification of reads. Similarly, skewed distribution of UMIs can lead to biased estimation of molecular counts. Finally, as droplet-based scRNA-seq protocols are still relatively new, detailed diagnostics and multiple quality control steps are typically needed to ensure high-quality measurements, and identify likely sources of problems. Given current lack of such general processing pipelines for the droplet-based scRNA-seq, we have set out to provide an open-source implementation.

## Results

We have developed a high-performance pipeline to perform initial pre-processing and analysis of the droplet-based scRNA-seq data. The pipeline characterizes the quality of a library using a wide range of diagnostic indicators, filters out artefactual cellular barcodes, evaluates and corrects for potentially confounding effects of uneven UMI coverage, and corrects for UMI and cellular barcode sequencing errors based on molecular similarity measures that do not require prior knowledge of the possible barcode sequences. It is designed to be used with different alignment methods and provides configuration options to accommodate alternative scRNA-seq protocol designs.

### Uneven UMI frequency distribution distorts molecular count estimation

In UMI-based protocols, the expression magnitude is typically estimated as a number of unique UMIs associated with a given gene in a given cell. If the space of possible UMI sequences is limited, it becomes possible for two separate molecules of the same transcript to be labeled by the same UMI. To account for such UMI collisions, Fu *et al*. have originally proposed a correction based on assumption of uniform distribution of UMIs in the overall dataset [3]. Such correction is rarely used, given relatively large numbers of possible UMIs. Examining droplet data from different protocols, however, we find that UMI frequency distribution tends to be highly skewed, with a small fraction of UMIs contributing to a disproportionately large number of molecules (Figure 1A,B, Figure S1). The outlier UMIs with the highest frequencies show aberrant mononucleotide stretches towards the end of their sequence. Such biases may arise to due errors generated during library construction protocol or truncated barcode constructs. Even when such erroneous UMIs are filtered out, the overall UMI distribution remains significantly skewed (Figure S1B,F), suggesting that a more advanced approach is needed to correct for the impact of UMI collisions. In implementing corrections for the UMI collisions, we therefore moved away from the assumption of a uniform UMI distribution, and used bootstrap procedure to directly model the true UMI frequency distribution (see Methods). This approach is effective at correcting UMI collisions on simulated data (Figure S2) and, as we will demonstrate in the next section, provides notable improvements on real data.

**Figure 1.**
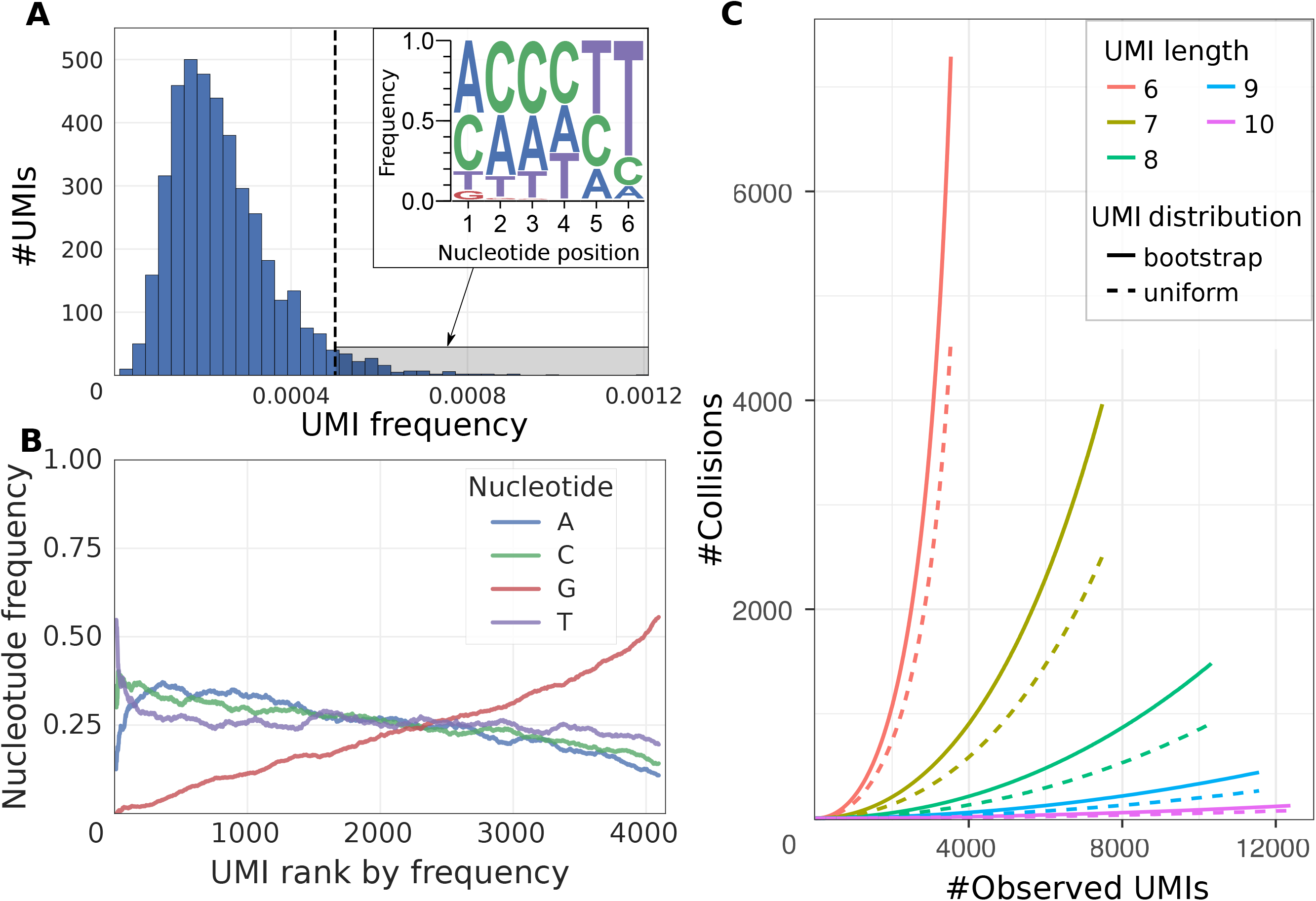
Skewed distribution of UMIs leads to increased number of UMI collisions. **(A)** Distribution of UMI occurrence frequencies across all genes is shown for the mouse ES dataset. The top-right inset shows position-specific nucleotide frequencies of the outlier UMIs (highlighted by gray shading on the main plot). Significant skewness of the UMI distribution decreases the effective pool of UMIs. **(B)** Proportions of different nucleotides in the UMI sequences are shown as a function of the overall uMi frequencies (x-axis orders UMIs so that most frequently occurring UMI sequences have low rank) for the mouse ES cells dataset. **(C)** Estimated number of UMI collisions as a function of the true gene expression level (x axis) is shown for different UMI lengths (simulated by trimming 10nt UMIs, see text). The estimates based on the uniform and empirical UMI distributions are shown. 10x Chromium human post-transplant BMMCs dataset was used. For short UMIs, the number of collisions observed at highly-expressed genes can be comparable to the true number of molecules. Longer UMIs decrease the number of collisions.

### Errors in UMI sequence lead to overestimation of molecular counts

An error introduced into UMI sequence during the library preparation can be mistakenly interpreted as an additional molecule. Computational corrections have been proposed to avoid such overestimation.

The simplest such approach [5] omitted for a given gene all UMIs that had an adjacent UMI sequence (Hamming distance equal to 1) with a larger number of reads (as in [6] we refer this method as *cluster)*. Indeed, the probability of having two molecules of the same transcript in the same cell being labeled by UMIs of Hamming distance 1 is low, given sufficient size of the UMI pool relative to the number of transcript molecules (see Methods). However, for moderately-expressed genes the observed number of such events exceeds the expected frequency by a factor of ~40 (Figure S3), suggesting that most adjacent UMI occurrences are erroneous. More complex, network-based solution [6] (referred here as *directional)* considers a UMI to be erroneous if it has an adjacent UMI with more than twice the number of reads.

An alternative approach, implemented in the 10X Chromium Cell Ranger pipeline [7], uses UMI base call quality to distinguish erroneous UMIs. Examining different droplet-based datasets, we find that the fraction of UMI errors that can be distinguished by lower base call quality varies between datasets, within the range of 29.4-85.6% (Figure S4). This suggests that a substantial fraction of UMI errors may originate during PCR amplification or other library preparation steps preceding the sequencing itself. Base call quality would not be informative in such cases. Furthermore, the existing methods do not consider the total number of molecules for a given gene, even though the probability of observing adjacent UMIs by chance increases. Such increase is further exacerbated by an uneven distribution of uMi frequencies described in the previous section. For instance, for a gene with 100 molecules and UMI length of 6nt, 36 adjacent UMIs would be expected under uniform distribution, and 40 under the empirically observed distribution (mouse BMCs dataset was used, see Methods).

To improve the accuracy of UMI filtering, we developed a Bayesian approach to estimate the posterior probability of a UMI being erroneous based on the gene expression magnitude, observed number of the adjacent UMI sequences, prior distribution of UMIs, as well as the position and identity of the nucleotide substitution (see Methods). To evaluate performance of different UMI correction approaches, we first examined a dataset with relatively long 10bp UMIs [8]. Given lower rate of accidentally observing UMIs at Hamming distance 1 in these longer UMIs, we used *cluster* UMI filtering procedure to obtain benchmark expression estimates for the dataset. We then simulated datasets with shorter UMIs by trimming the UMI sequences, and comparing the resulting molecular count estimates to the corresponding full-length benchmark values (see Methods).

While errors in the UMI sequences lead to over-estimation of the molecular abundance, UMI collisions lead to under-estimation. The probability of such collisions increases for shorter UMIs, which results in pronounced under-estimation of molecular counts at short UMIs (Figure 2A). Conversely, the overestimation due to sequencing errors is more apparent at longer UMIs. Comparing different UMI collision correction methods, we find that the proposed approach based on bootstrap sampling of the empirical UMI frequency distribution shows much better performance than correction based on the uniform UMI distribution assumption (Figure 2B). We then compared different methods for correcting UMI sequence errors. In addition to the standard *cluster* algorithm [5], we also evaluated a variant that disallows merging of UMIs of equal sizes *(cluster-neq)*. We found that the Bayesian approach proposed here significantly outperforms existing methods (Figures 2C, S5, Tables S1-4). The impact of both collision and sequencing error corrections is most notable for genes within the high expression range.

**Figure 2.**
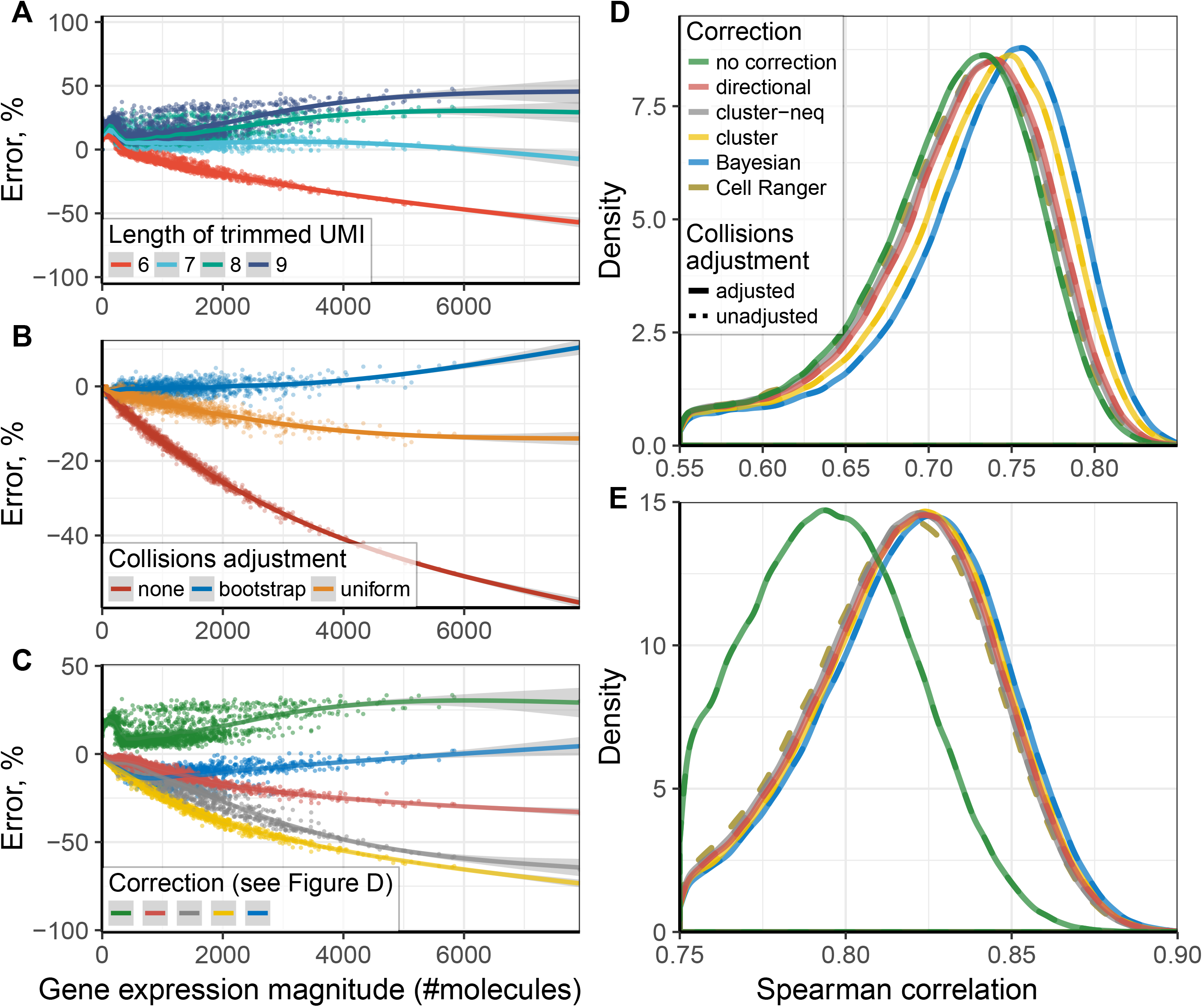
Comparison of UMI collision and sequencing error correction methods. **(A)** The scatter plot shows percent error (y axis) in estimation of the molecular counts for different genes using computationally trimmed UMIs (down to 6-9nt lengths) from their original 10nt length, as a function of the full-length UMI estimate (x axis, see Methods). The line shows spline-smoothed dependency with the 95% confidence band. The errors result from two opposing trends, with UMI sequencing errors inflating the resulting count estimates, and UMI collisions deflating the estimates. Shortened UMIs result in larger number of collisions. **(B)** The effect of different UMI collision corrections is shown on the 6nt trimmed UMIs. **(C)** Comparison of different UMI sequence error correction methods is shown for the 8nt trimmed UMIs. UMi collisions were corrected using *bootstrap* approach in all cases except for “no correction”. **(D, E)** The impact of UMI corrections on the expression cohesion within the resulting cell subpopulations. Distribution of within-cluster expression Spearman rank correlations is shown for the human BMMC dataset (D), and human PBMC dataset (E). UMI errors and collisions decrease within-cluster similarity (assessed here using transcriptome-wide Spearman rank correlation). Improvements resulting from different UMI error correction methods are shown, with (solid) or without (dashed) bootstrap collision adjustments.

To examine the impact of different approaches on the downstream analysis we evaluated the similarity of expression profiles within major cell subpopulations in datasets generated using inDrop [9] and 10x Chromium [10, H] platforms. We found that the proposed corrections can result in significantly higher expression profile correlation of cells belonging to the same cluster, compared to uncorrected molecular counts (Figure 2D, S7). To ensure robustness of the comparison we used pairwise Spearman rank correlation, which was estimated over the genes expressed in both cells of the pair. The proposed Bayesian approach showed best performance, though the difference in correlation gain between different correction methods was small in some datasets (Figure S7).

### Correction of the cellular barcode sequence errors

The number of different cellular barcodes (CBs) in a droplet-based library normally exceeds the number of actual encapsulated cells by several fold (Figure 3A). Similar to issues encountered in UMIs, additional CBs can result from sequence errors introduced during library construction or sequencing. This would result in material from one droplet being mistakenly split up into several different CBs. Alternatively, additional CBs may also be empty droplets that did not encapsulate a real cell, but instead captured background RNA or cell debris together with an indexing bead [12]. To evaluate whether this is a significant factor, we examined the read composition in an inDrop control experiment which encapsulated a mixture of human and mouse cells (see Methods). We find that the fraction of reads originating from the admixed genome increases with decreasing cell size (Figure 3B). Furthermore, we find that the admixture fraction correlates positively with the mitochondrial read fraction. These results suggest that capture of extracellular RNA background or cell debris by empty droplets provides noticeable contribution to small CBs.

**Figure 3.**
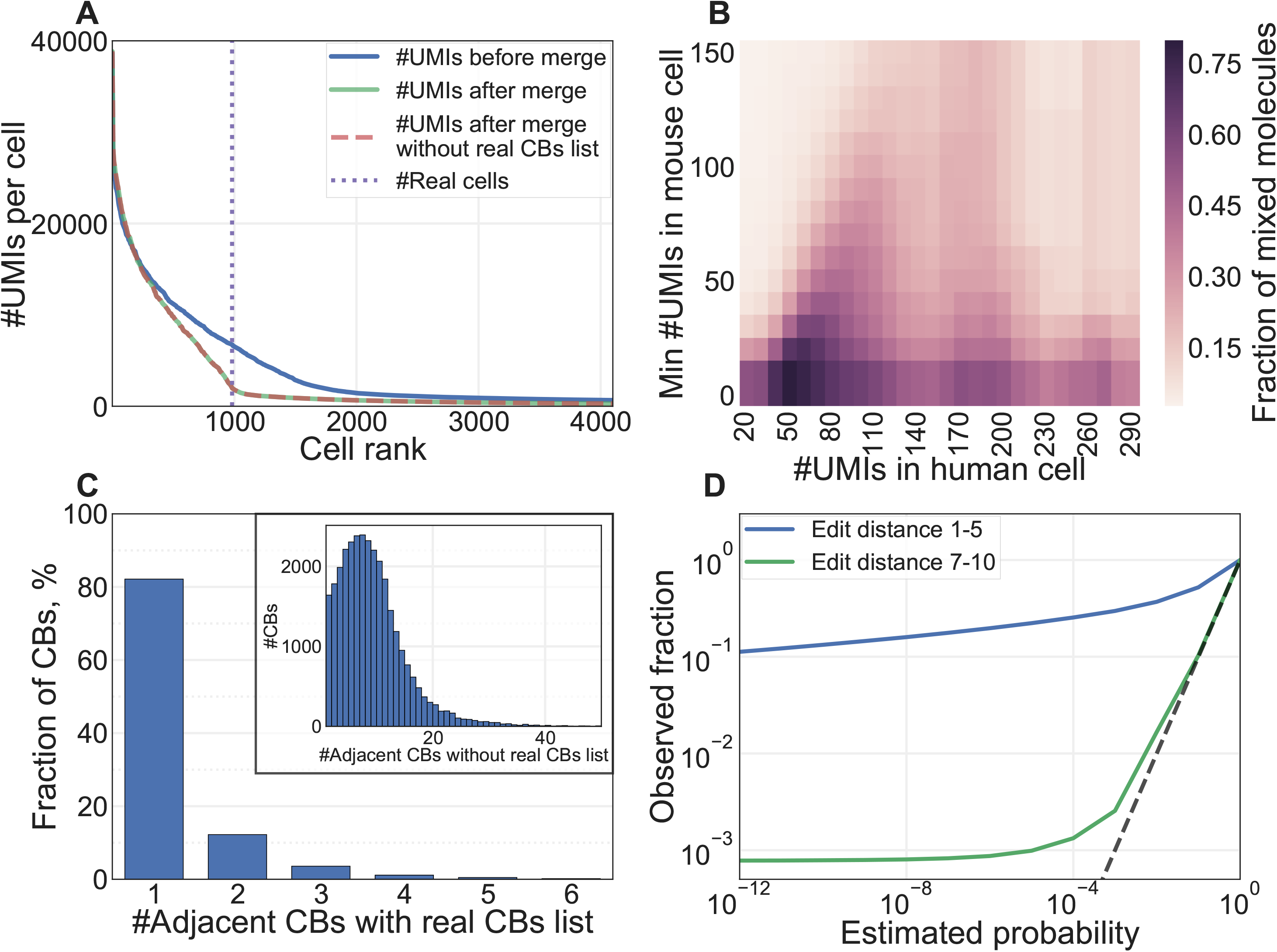
Correcting for Cellular Barcode errors. **(A)** The number of molecules per cell before and after the merge correction procedure, is shown for the mouse ES dataset as a function of the overall cell size as reflected by rank of the cell (with cells possessing most molecules having the lowest ran; cells with ≤ *500* molecules were omitted). The results of the merge procedure with and without the prior knowledge of the possible barcode list match almost exactly. The resulting distribution shows a pronounced inflection point coinciding with the real number of encapsulated cells [1]. **(B)** Analysis of human-mouse cell mixtures suggest contribution of extracellular background. The fraction of human-mouse molecule admixture is shown (color gradient) for cells binned by the number of molecules originating from the two genomes. The cells were binned by the number of human molecules (x axis), and the fraction of mouse molecules originating from mouse cells exceeding a certain size threshold (y axis) was calculated for each bin. The majority of admixed molecules originate from small mouse cells, with highest admixture fraction observed between small mouse and small human cells (>75%). **(C)** The number of equidistant adjacent CBs of larger size is shown for each of the observed CBs in the mouse ES dataset. The main plot shows adjacent CBs selected from an *a priori* known set of valid CB sequences. The inset shows counts of adjacent CBs selected from all CB sequences observed in the dataset. **(D)** Comparing all pairs of cells with similar CBs (edit distance 1-5) and distant CBs (distance 7-10), the plot shows the dependency between the theoretical probability of a certain number of overlapping UMI-gene combinations between the two cells (x axis) with the observed empirical frequency (observed fraction, y axis) of such cells. Cell pairs with similar CBs show much higher overlap, which is driven by CB sequencing errors. A small fraction of distant CBs (0.1%) also shows higher molecular overlap than expected, which is likely explained by cross-droplet contamination.

We first examined methods for correcting CB sequence errors. In doing so we considered two scenarios: one where a list of possible valid CB sequences is known (e.g. 10x or inDrop), and another where CBs can be an arbitrary nucleotide sequence (e.g. Drop-seq). Pre-designed CB sequences are typically evenly spaced in the sequence space, and replacing an erroneous CB with the closest matching valid CB sequence is an effective strategy. The space of potential valid CBs can be further narrowed down by taking into account that the valid Cb shouldn’t have fewer counts than the erroneous CB. However, if the list of possible valid CBs is unknown, or if there are many similar CBs *(e.g*. short barcodes), the number of possible merge targets increases significantly (Figure 3C). To accurately determine the probability that two CBs originated from one CB, we used UMI-gene composition similarity, which evaluates the likelihood that two independent cells will end up producing equivalent UMI-gene combinations (see Methods). We evaluated the model on the mouse ES dataset by estimating the intersection probabilities for the cells with distant CBs (Hamming distance ≥7), which were unlikely to be split up due to CB sequencing errors (Figure 3D). Indeed, it can be seen that almost all UMI-gene intersections between such cells could be explained by chance occurrence. Approximately 0.1% of all pairs of cells with distant barcodes show highly significant degree of molecular overlap (p<10^-5^), which is likely caused by another error modality such as RNA background contribution described above. To validate precision of the implemented merge procedure we compared results obtained with and without the usage of the real CB list. When analyzing mouse ES dataset without the knowledge of valid CBs, the algorithm performed 97.6% of the same merges into the real CB sequences resulting in nearly identical set of top cells (Figure 3A). Thus, the implemented approach provides effective correction for CB errors even when the set of possible CB sequences is not known, such as Drop-seq measurements.

### Recognizing damaged or low quality cells

The number of molecules associated with a given CB generally provides reasonable criteria for selecting real cells [1, 7]. Similarly, CBs with very few associated reads likely represent empty droplets. However, classifying CBs in the intermediate range poses a challenge. The intermediate size CBs likely contain damaged or dying cells from which relatively little mRNA material could be recovered [12]. This complicates the optimization of a size separation threshold. Such low-quality cells could also cover a range of sizes, making the use of a single size cutoff ineffective.

Classification of low quality cells was examined by Illicic *et al* [12], where an support vector machine (SVM) classifier was trained based on examination of cell morphology from microscopy data prior to lysis and library preparation. As such data is difficult to obtain for the droplet approaches, and existing SVM cannot be directly applied to different protocols or cell types, we aimed to develop a self-contained approach that would not require high-quality training data. While the true labels for low-and high-quality cells are not available, we argued that large cells initially include a large fraction of high-quality cells, and small cells include very low fraction of high-quality cells. We then aimed to train a classifier to distinguish high-quality cells based on a limited set of technical features (see Methods), taking into account that the initial labels of the training set will contain some fraction of errors. The tolerance of different classifiers to training set errors can vary considerably. We evaluated performance of several appropriate approaches (KDE [13], Random Forest [14] and Robust Gaussian Processes [15], see Methods). In addition to cross-validation score we measured robustness of the classifiers with respect to: removal of a random 20% of the training data (5-fold cross-validation, see Table 1); introduction of artificial noise into the data (Figure S9A,B); narrowing/widening of the thresholds used to separate large and small cells for the initial label assignment (Figure S9C). Based on the resulting performance and runtime complexity *(e.g*. Robust GP has a high complexity of O(n^3^) relative to the number of samples) we chose Kernel Density Estimation (KDE) classifier. An example of KDE-based quality scores is shown in Figure 4A for the mouse pancreatic dataset. While the quality scores show expected association with cell size, some of the smaller cells are able to attain high scores, and some of the large cells are assigned low scores (Figure S10, S11). The additional “high-quality” (score>0.9) cells identified by the classifier beyond the initial size threshold show consistent clustering with the major subpopulations (Figure 4B) supporting their validity. In contrast, selecting a set of cells on the basis of the size threshold alone quickly leads to appearance of an artefactual cluster of poor-quality cells (Figure S12). Filtering of cells by the KDE quality score preserves all of the subpopulations identified by the original publication [9]. An example of cell classification on another high-complexity dataset (mouse bone marrow), shown in Figure 4C also illustrates poor clustering of cells with low quality scores.

**Figure 4.**
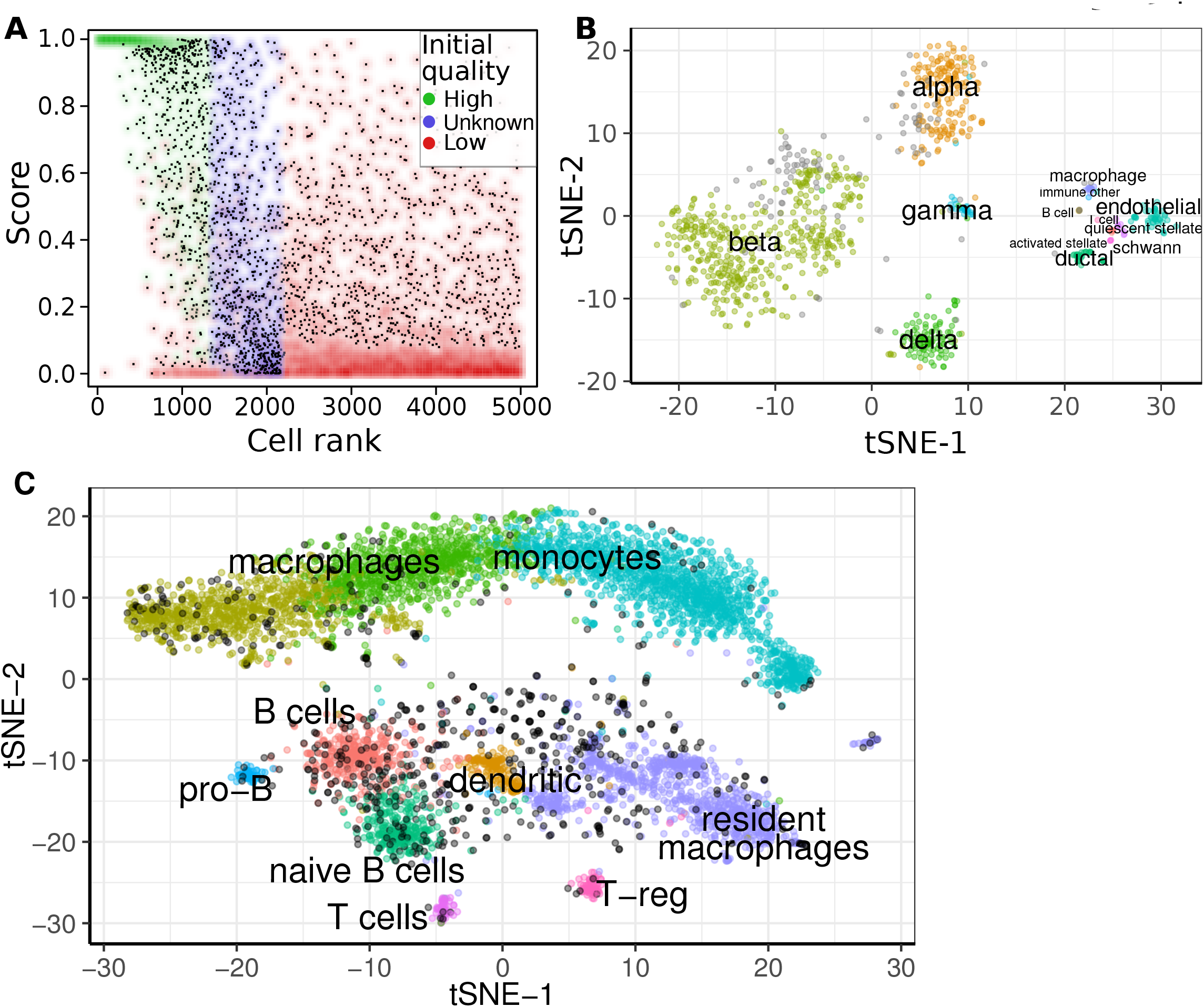
Classification of low-and high-quality cells. **(A)** The cell quality score (y axis) determined by the developed approach is shown as a function of cell size (x axis, shown as rank with largest cells being assigned the lowest rank) for the mouse pancreatic cell dataset paper [9]. The initial size-group assignment of each cell is shown by color. As expected, most large cells are reported to have high scores, and cells with small number of molecules tend to have low quality scores. Nevertheless, some of the large cells are reported as being low in quality and some of the small cells are reported to have high quality. **(B)** Visualization of high-quality cells (score>0.9) using t-SNE projection in mouse pancreatic dataset. The cells are colored according to the original cluster labels, with cells that were not passing the original size threshold but were nevertheless deemed to be of high quality shown in gray. Most of such cells can be confidently assigned to the annotated clusters. **(C)** Visualization of low-quality cells using t-SNE projection on mouse bone marrow dataset. All cells passing the initial size selection threshold are shown (see Methods). Cells with quality scores below 0. 9 are shown in black. These poorly-scoring cells are generally found scattered outside of the dendritic and resident macrophage clusters.

## Discussion

Droplet-based microfluidics protocols and other high-throughput methods are enabling production of large single-cell RNA-seq datasets (10^3^-10^6^ cells). Complex barcoding schemes employed by such methods require in-depth computational analysis to achieve accurate recovery of molecules associated with different cells and genes. In order to avoid collisions of cellular barcodes, large numbers of cells necessitate longer CBs, increasing the probability that a sequence error will be introduced into a CB during the bead construction steps [1], library preparation procedures, or library sequencing. We show that in many such errors there are multiple equidistant CBs from which the molecule may have originated. The implemented solution, which merges CBs based on the probabilistic assessment of the molecular overlap between the CBs, provides accurate correction even in the cases when the set of possible valid CBs is not known in advance.

Errors affecting molecular barcodes (UMIs) pose a similar challenge, which in this case is driven by increasing sequencing depth of individual cells. This has been recognized by earlier studies [5], and several correction strategies have been proposed. We show that the overall distribution of UMI sequence occurrences is not uniform, and the resulting bias reduces the effective UMI space leading to increased number of UMI collisions in well-expressed genes and deflated molecular counts. Some of the UMI errors appear to result from occurrence of aberrant library molecules incorporating mononucleotide primers, such as polyT into the UMI position. On the other hand, point mutations in UMIs and aberrant base calls can lead to inflated molecular counts. While most UMI errors can be mitigated experimentally by increasing the UMI length, we show that taking into account empirical distribution of UMI frequencies allows to adjust for both UMI collision and sequence error effects.

Even with corrections of CB sequence errors, most of the CBs encountered in the current droplet-based datasets do not represent real cells. These additional molecules may originate from empty droplets capturing extracellular background. Indeed, examination of mouse-human dataset mixtures, suggests that smaller CBs have higher cross-organism contamination fraction that one would expect from extracellular background. In addition to empty droplets, some of the low-magnitude CBs may represent damaged, dying or dead cells, as well as cells that were not successfully measured for other reasons. The challenge of identifying damaged cells has been previously examined by Ilicic *et al*. [12] in the context of Fluidigm C1 protocol where the proportion of low-quality cells is typically in the range of 10-40%. This fraction can be much higher in the inDrop data *(e.g*. 90% of CBs), and obtaining microscopy-based labeling for the classification is challenging given the rapid flow within the devices. We instead, explored application of fault-tolerant classifiers to identify technical features consistent with an imperfect initial separation of high-quality cells based on the size criteria alone. Such approach is able to pick up relatively large cells that resemble poorly-measured cells based on their technical features, and rescue some of the smaller cells that look consistent with high-quality tail of the cell distribution. Overall, we hope that the developed pipeline will facilitate analysis of the droplet-based single-cell RNA-seq data, providing helpful diagnostic (see Supplementary Note 1 for an example of a dropEst pipeline report), and improving the accuracy of the resulting expression estimates.

## Methods

The dropEst pipeline operates in three phases: i) identifier parsing phase; ii) read mapping phase; and iii) filtering and quality control phase. The first phase takes as an input FASTQ files containing paired-end read and index data. The output of this phase is a modified FASTQ file with reads which can be aligned to a transcriptome reference during the second phase using a standard splice-aware aligner *(e.g*. STAR [16], or Tophat 2.1.0 [17] that was used in our work). The third phase takes BAM files with the aligned reads [18] and a gene annotation file in GTF format. BAM files produced by the 10x Cell Ranger pipeline can be also provided when running this stage. The result of pipeline is an R-readable file that contains molecular count matrix and other processed information, as well as a report with diagnostic information on the library.

### Correction of UMI collisions

In case when the number of UMI per gene is comparable to the total UMIs pool size, the gene expression level will be underestimated [3], and needs to be adjusted by taking UMI collisions into account. Fu *et al*. [3] assumed uniform distribution of UMI probabilities and the number of unique UMIs expected for number of molecules *n* from a UMI pool of size 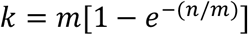, thus 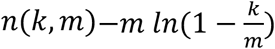. To account for non-uniform UMI distribution observed in the droplet datasets, *n(k,m)* was estimated computationally using UMI bootstrap sampling. Specifically, possible values of *k* were quantized with a step value 50. For each quantized value of *k*, a 1000 bootstrap sampling iterations were performed to estimate corresponding values of *n*. Subsequently, in estimating *n(k,m)* for an arbitrary value of *k*, linear interpolation was used. To validate the developed method, we simulated UMI collisions by randomly merging small genes (Figure S2).

### Correction of UMI sequence errors

To determine whether two UMIs represent technical variations of the same UMI, we use Bayesian approach to derive an optimal decision boundary based on the overall UMI frequency distribution.

Given two UMIs within a gene, we considered the following features:

- *U:* sequence of the first UMI.
- *u:* sequence of the second UMI.
- *R:* number of reads for the first UMI.
- *r:* number of reads for the second UMI, *r ≤R*.
- *N_S_:* number of adjacent (Hamming distance of 1) UMIs for the UMI *U* with the number of reads *r^'^: r_'_ ≤ R*.
- *N_L_:* number of adjacent UMIs for the UMI *U* with the number of reads *R^'^:R^'^ > R*.
- *S_g_:* number of UMIs in the gene.

in the following description, when referring to the UMI with the larger number of reads we use the upper case latters, and for the UMI with the fewer reads we use lower case letters. Let 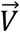 denote *(U, R, N_s_, N_l_)*. The equation of the optimal decision surface:

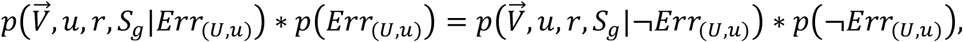

where *Err_(U,u)_* means that *u* occurred because of an error in *U*.

As we have no information about prior distribution of the events *Err_(U,u)_)* and *¬Err_(U,u)_*, we assume them to be equally likely:

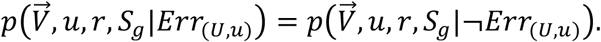

The right part of the equation can be simplified as follows:

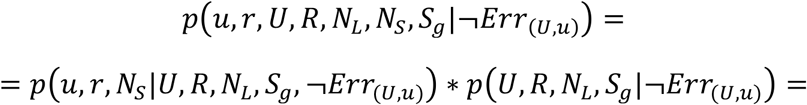

Events *U, N_L_*, do not depend on the *Err_(U,u)_* (let's suppose the decrease in *R* insignificant). Also we assume that *S_g_* does not depend on the *Err_(U,u)_*. Then the right part of the equation takes the form:

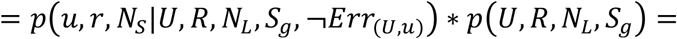

Given *¬Err(_U,u)_*, events *u* and *r* do not depend on 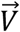:

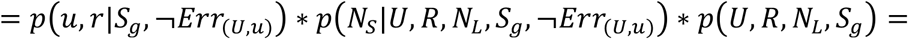

Given *Err(_U,u)_*, event *N_L_* and *N_S_* does not depend on *u*, and *r* does not depend on *S_g_*. Furthermore, *¬Err_(U,u)_* does not mean that *u* is a real UMI, as *u* could be a result of an error in a UMI other than *U*. Then *p(r|¬Err_(U,u)_) ≈ p(r)*. Similarly, *p(u|¬Err_(U,u)_) ≈ p(u)*:

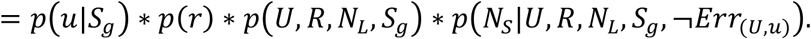

The left part of the equation could be simplified as follows:

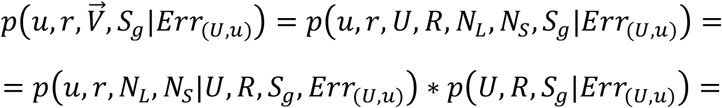

Given *Err_(U,u)_*, events *N_L_* and *N_S_* depend on *u*, and on *r* only through *U* and *R*. Moreover, *R* does not depend on *U* and *S_g_*, and events *U* and *S_g_* do not depend on *Err_(U,u)_*:

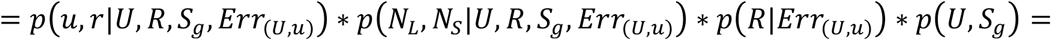

Given *Err_(U,u)_*, event *u* depends on *U* only through the position of the difference between the sequences *C_(U,u)_* and the nucleotide replacement *D_(U,u)_*. Also, neither *u* nor *r* depends on *S_g_*:

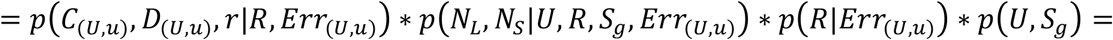

Events *C_(U,u)_* and *D_(U,u)_* do not depend on *r* and *R*. Also, we can assume them to be independent from each other:

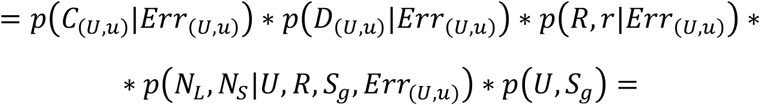

We denote 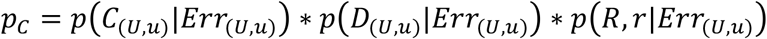:

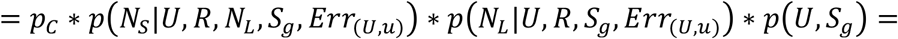

*N_L_* doesn't depend on *Err_(U,u)_*:

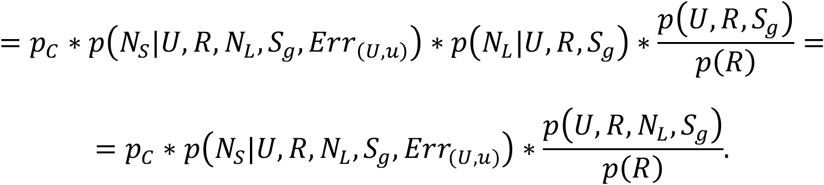

The simplified expressions can be combined as follows. The equation of the Bayes optimal decision boundary can be written as:

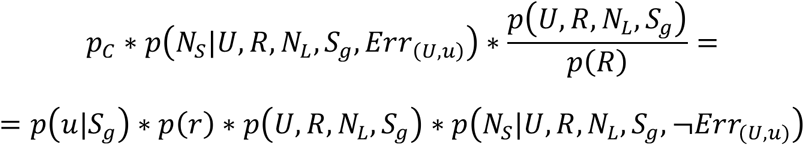

or

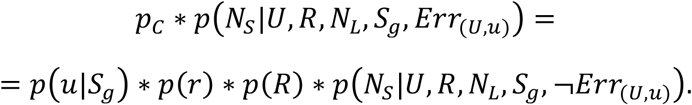

After reordering:

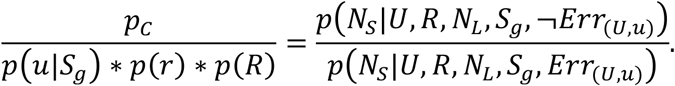

As *p(u|S_g_ = (1-(1-p(u))^S_g_^*):

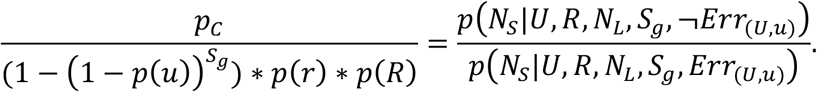

Let's suggest that *N_s_* depends on *R* only through *N_L_*:

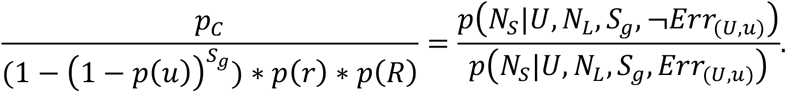

### Estimating probabilities

*p(u), p(r)* and *p(R)* can be simply estimated by the observed frequencies. To estimate probabilities *p(C_(U,u)_|Err_(U,u)_)*, *p(D_(U,u)_|Err_(U,u)_)* and *p(R,r|Err_(U,u)_)* we create a training sample, which contains only pairs of UMIs where *u* occurred because of an error in *U*. Such set was assembled by choosing the genes containing only two adjacent UMIs only. The theoretical probability *p(u, U|S_g_ = 2, ¬Err_(U,u)_* is negligible. Thus we expect almost all such events have occurred because of an error in *U*. Such training sample is representative because all events *C_(U,u)_*, *D_(U,u)_*, *R* and *r* are independent of *S_g_*. Thus, estimation of *p(C_(U,u)_|Err_(U,u)_)* and *p(D_(U,u)_|Err_(U,u)_)* becomes straightforward. Probability *p(R,r|Err_(U,u)_)* was approximated by the multinominal distribution.

Estimation of 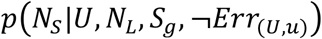 and 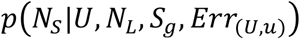. These probabilities depends on the large numbers of parameters, making the training approach impractical. We use theoretical estimation of 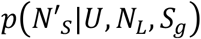, where 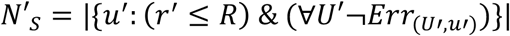 (see the algorithm below). To simplify theoretical estimation, we can use the following procedure. We introduce a scoring function for the pair of UMIs 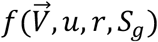, which is approximately proportional to the probability 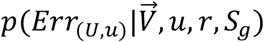. We then assume that the event *¬Err_(U,u)_* is equal to the event 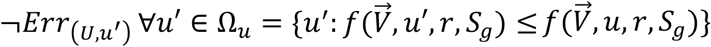 Similarly, *Err_(U,u)_* is equal to the event 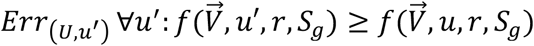 Then we estimate:

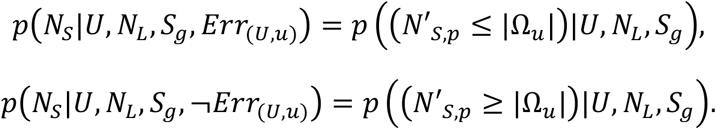

We can use the left part of the decision boundary equation as a scoring function 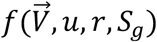. In this case, for the fixed UMI *U* within the fixed gene *g* we can order all of the adjacent UMIs *u_i_* by the increasing of 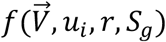. The decision boundary equation:

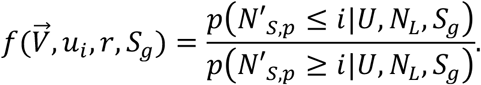

Estimating distribution ***p(N_'s_|S_g_, U,N_L_)***.

We use the following notations:

- *L:* length of the UMI.
- *N_UMI_:* total number of possible UMIs (in most cases is equal to *4^L^*).
- *K:* maximal number of the adjacent UMIs (in most cases is equal to *3L)*.
- *P_Neighb_ = P_Neighb_(U)*: probability to observe a UMI, adjacent to *U*. Equal to *Σ_uϵAdjacent(U)P(u)_*.
- *N'*: total number of adjacent UMIs for the UMI *U*.

To estimate the distribution of *N^'^_s_* we use the following assumption:

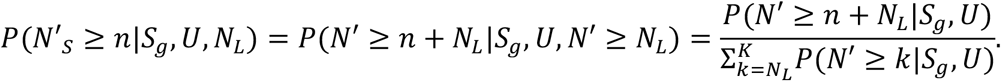

The distribution *p(N^'^|S_g_,U)* was estimated by modeling of the process of picking UMIs from the pool. Suppose that we already picked *s* UMIs and we have *k* different adjacent UMIs. Let us denote it as the state (*k,s*). This state can occur in of one of the following situations:

1. We were previously in a state *(k,s - 1)* and picked a UMI which was not a new adjacent UMI (*i.e*. either a previously observed adjacent UMI, or not adjacent UMI). The probability of such pick is 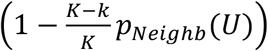.
2. We were previously in a state *(k - 1,s - 1)* and picked a UMI, which is a new adjacent UMI. The probability of such pick is 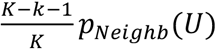.

The model above can be evaluated using dynamic programming. To do so we build a matrix *T = {t_k,s_}*, each cell of which contains the weighted sum of the neighboring bottom-left and left cells in the matrix T (see example in Table 2).

Such matrices would need to be computed for the each UMI present in the dataset. However, the asymptotic of this approach is equal to *0(S_g_ * #UMI * K)* in terms of both time and memory, which would be too large for large datasets. To optimize it we employ the following solution. The matrix *T* depends on *U* only through *p_Neighb_(U)*, and the rate of change of the function within a cell is proportional to *p_Neighb_(U) + o(p_Neighb_(U))*. Thus, we assume *p(N^'^|S_g_, U)* to be a piecewise constant function from *P_Neighb_(U)* and perform a quantization by this probability. A quantization step *Δp = 0.01* was used.

Iterative procedure of UMI sequence errors correction. After the estimation of the decision boundary, all UMIs that were determined as erroneous are removed. This will change the input parameters *S_g_,N_L_* and *N_s_* of the algorithm. Therefore, to perform a precise filtration, the procedure is ran iteratively.This does not add significant amount of runtime complexity because: *i)* only the right part of the decision rule is affected; *ii)* dynamic programming matrices are calculatd only once, as the gene size cannot increase during filtration; *iii)* for genes with a small number of UMIs, the procedure converges after one or two iterations.

### Correction of cellular barcode sequence errors

CB sequence errors split a fraction of the molecules originating from one cell into smaller CBs. Given that the number of reads per UMI is generally high then one, the smaller CBs will contain some of the same gene-UMI combinations as the true CB. In other words, the smaller CBs will have similar molecular composition - the set of cell unique genes-UMI combinations. We use composition similarity as a criterion for determining whether the two barcodes should be merged. Size of the compositional intersection between two independent cells is modeled using Poisson distribution with the mean dependent on the UMIs distribution and the number of molecules associated with the CBs. For the two fixed CBs with molecular count vectors (gene expression profiles) 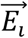 and 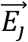 the intensity parameter 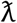 of the Poisson distribution was estimated using the following bootstrap-based procedure:

1. The distribution of UMI probabilities *P_0_* was estimated over all genes in all cells of the dataset.
2. Two sets of UMIs with sizes 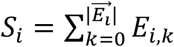 and 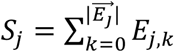 were taken from *P_0_* with replacement.
3. The selected sets were randomly allocated to genes, according to the profiles 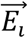 and 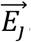.
4. The number of common gene-UMI combinations was counted.
5. The procedure was repeated a fixed number of iterations, starting with the step 2.

Using the estimated distribution, we perform a statistical test for the hypothesis *H_0_:* observed size of the intersection *S_*intersection_* was obtained by chance. P-value of the test is the value of the Poisson distribution function 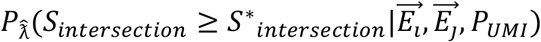, where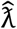 is the estimated parameter.

The implemented pipeline uses this test to compare each cell *C_i_* with all cells *C_j_* that have higher total number of molecules *(S_j_ ≥ S_i_)* and whose CBs have Hamming distance lower than a fixed constant. In the presented results the constant was taken to be 2. To account for multiple comparisons, we used Bonferroni correction.

### Classifying damaged or low-quality cells

The implemented approach for classification of damaged and low quality cells split into three tasks: (i) creation of the training sample, *i.e*. establishing the initial class labeling, (ii) feature selection and (iii) application of a classifier algorithm.

The initial class labels were assigned based on the cell size. To do so, the dataset was split into three parts: ‘big’ cells, ‘intermediate’ cells and ‘small’ cells. To determine the borders of ‘big’ and ‘small’ cells we used the plot log `(#UMI in cell) vs. log (cell rank)` (Figure S8C). This heuristic is based on an observation that the left part of such plot has a negative second derivative, followed by a linear part, and a third part of the plot has a positive second derivative. The automated procedure locates the upper *(t_u_)* and the lower *(t_l_)* position of the linear part. The cells with size smaller than *t_l_* were assigned the initial label of ‘low-quality’ cells. The top 75% of the cells with the size larger than *t_u_* were assigned the initial label of ‘high-quality’ cells. The remaining cells were labeled as ‘unknown’. Alternatively, the initial label borders can be specified by hand, for instance based on the shape of the log `(#UMI in cell * #Cells) vs log (#UMI in cell)` plots (Figure S8A,B).

Given the initial labeling, the cell classification problem was considered as a problem of establishing robust classification in the presence of training label noise [19]. We compared three classifiers: Kernel Density Estimation classifier [13], Random Forest [14] and Robust Gaussian Processes Classifier [15]. Following evaluation of robustness we chose KDE classifier with Normal Scale bandwidth selector [20] (the implementation provided by the R package ‘ks’ was utilized [21]).

Two types of features could be considered: biological features *(e.g*. expression levels of genes belonging to different GO categories [22, 23]), and technical features *(e.g*. different statistics on the sequenced data). We expect most biological features to be dataset and cell type-specific [12], with exception of the mitochondrial fraction, which has appeared as a robust indicator of cell death across most datasets [12]. Therefore, in choosing classifier features we limited biological features to the fraction of UMIs on mitochondrial reads. The following technical features were utilized:

1. Mean number of reads per UMI.
2. Mean number of UMI per gene.
3. Fraction of low-expressed genes (genes with 1 molecule).
4. Fraction of UMIs on low-expressed genes.
5. Fraction of intergenic reads.
6. Fraction of not-aligned reads (optional feature, as it typically has to be calculated during the identifier parse phase).

The computational complexity of the KDE classifier estimation has exponential dependency on the dimensionality of the feature space. We therefore reduced the feature space by using the first three principal components of the feature space for classification. Additional constraints on the mitochondrial fraction were applied prior to classification: all cells with high mitochondrion fraction are explicitly set as ‘low-quality’ prior to the algorithm training. High fraction threshold was set to be *m+4a*, where *m* is the median cell mitochondrial fraction in the dataset, and *a* is the median absolute deviation of the mitochondrial fraction.

### Mouse Bone Marrow inDrop measurements

Whole bone marrow cells were isolated from 11 weeks old C57Bl/6 male mice (Jackson Laboratory). Epiphysis/metaphysis fraction from long bones was collected, crushed, cut into small pieces, and digested using Collagenase I (STEMCELL Technologies) for 30 minutes at 37-degree agitation. Bone marrow cells were filtered through a 70-micron filter. Red blood cells were lysed using Ack-lysis (ThermoFisher Scientific) on ice for five min, quenched with Media 199 (ThermoFisher Scientific) supplemented with 2% fetal bovine serum (ThermoFisher Scientific), and spun down at 500 g for 5 min. Cells were stained for 30 minutes with the red blood cell marker TER119 (Biolegend) and cells were sorted using DAPI (ThermoFisher Scientific) as a live/dead viability marker. Four hundred thousand live whole bone marrow cells (negative for TER119) were sorted into medium 199 (ThermoFisher Scientific). Before inDrop encapsulation cells were counted using a Cellometer (Nexcelom Bioscience). Cell viability is over 90%.

InDrop processing. Concentration of cells was adjusted to 300,000 cells/ml by adding PBS to the sorted cells. The cell suspension was then mixed 1:1 (v/v) with PBS containing 30% OptiPrep Density Gradient Medium (Sigma D1556) to obtain 150,000 cells/ml in 15% Optiprep. Using four microfluidics pumps and a polydimethylsiloxane (PDMS) microfluidic device, about 10,000 cells were co-encapsulated with barcoded polyacrylamide beads and a reverse transcription mixture containing Superscript III into water-in-oil droplets, according to a published protocol [24]. The library preparation and quality control procedures were carried out as described [24]. Indexed libraries were pooled and sequenced on aNext-seq 500 system (Illumina) at 2 pM concentrations.

### Mouse-human cell line mixture inDrop measurement

CK1750 mouse lung cancer cells (Carla Kim laboratory, Boston Children's Hospital) and K562 human immortalized myelogenous leukemia cells (ATCC) were mixed at 1:1 ratio to obtain 70,000 cells/ml in PBS containing 15% Optiprep. About 3,000 cells were co-encapsulated with barcoded polyacrylamide beads and a reverse transcription mixture containing Superscript III into water-in-oil droplets; and a library was prepared according to a published protocol [24]. The library was sequenced on a MiSeq system (Illumina).

### Datasets

The following datasets were used in evaluating the developed pipeline:

1. Mouse embryonic stem cells (935 cells, SRA accession SRR1784310 (GEO GSE65525), [1])
2. Human pancreatic cells (sample 4, run 2, SRA accession SRR3879612 GEO (GSM2230760), [9])
3. Mouse pancreatic cells (sample 2, SRA accession SRR3879617, SRR3879618, SRR3879619 (GSM2230762), [9])
4. Mixture of human and mouse cells (2876 cells, SCG29, GEO GSEXXXX)
5. Human frozen peripheral blood mononuclear cells (2900 cells, 10x Genomics Frozen PBMCs (Donor A), [7, 10])
6. Human pre-transplant bone marrow mononuclear cells (900 cells, 10x Genomics AML035 Pretransplant BMMCs, [7, 25])
7. Human post-transplant bone marrow mononuclear cells (900 cells, 10x Genomics AML035 Posttransplant BMMCs, [7, 8])
8. Human frozen bone marrow mononuclear cells (2400 cells, 10x Genomics Frozen BMMCs (Healthy Control 2), [7, 11])
9. Human CD34+ cells (9000 cells, 10x Genomics CD34+, [7, 26])
10. Mouse bone marrow cells (SCG71, GEO GSEXXXX)

### Implementation

The dropEst pipeline implementation is available on github:

## Acknowledgements

We would like to thank Allon Klein and Sinisa Hrvatin for their helpful comments on analysis of inDrop datasets, Carla Kim for providing mouse cell line for inDrop testing. PVK was supported by NIH R01HL131768 from NHLBI.

The authors declare no competing interests in regards to this manuscript.

## Supplementary Figure Legends

**Figure S1. Skewness of UMI distributions.**

As in Figure 1 of the main manuscript, nucleotide frequencies and UMI distributions are shown for mouse ES cells dataset after the filtration of extreme right tail (A, B). The analogous plots are also shown for 10x the human post-transplant BMMCs dataset before (C, D) and after the filtration (E-F). The filtering only has a visible effect on the left peak of the distribution, and does not affect the overall skewed shape of the distribution.

**Figure S2. Simulation of UMI collision frequencies.**

To evaluate the developed approach for correcting UMI collisions, we modeled collisions by randomly merging genes within a given cell (using mouse BMC dataset). The number of the resulting unique UMIs after the merge is shown on the x axis. The y axis shows the number of collisions *(i.e*. the difference between the total number of unique molecules before and after the merge). The observed number of collisions in different genes is shown in red dots. The collision frequencies predicted by different models are shown by lines. Bootstrap sampling from the observed distribution shows good fit to the data.

**Figure S3. Probability of observing adjacent UMIs in small genes.**

Probability of observing two adjacent uMls in a given cell in a given gene is shown for different edit distances (x axis). Probability estimated based on the bootstrap procedure is shown in blue. The empirically observed probabilities are shown in green. Mouse ES dataset was used. The observed number of adjacent UMIs exceeds the expected estimates by a factor of 40, indicating that most of the adjacent UMI occurrences result from UMI sequence errors.

**Figure S4. Recognition of UMI errors by base calling quality.**

Errors in UMI sequence can occur during either amplification or sequencing. Depends on the source, these errors can or cannot be distinguished by base calling quality. To examine the contribution of these mechanisms to UMI sequence errors we compared distributions of base call quality by different subsets of data. We focused on the adjacent UMIs (same gene, same cell), most of which are expected to be erroneous (Figure S3). The boxplots show for each dataset, the distribution of nucleotide base call quality for the adjacent (red) and distant (blue) UMIs. Specifically, the red box plots show base call quality for the nucleotide discrepant between the adjacent UMIs. The blue box plots show quality distribution across all nucleotides in pairs of UMIs that are sufficiently distant (Hamming distance > 3). The numbers above the box plots show the fraction of reads of each type that were correctly classified based on the base call quality alone (using naive Bayes classifier). The results suggest that the fraction of UMI errors that can be easily explained by the base call quality varies between the datasets, ranging from 29.4% to 85.6%.

**Figure S5. UMI sequence correction comparisons on trimmed UMI data.**

**(A)** Boxplots show distributions of errors (expressed as #UMIs+1 on a log scale) for different datasets, for different lengths of the trimmed UMIs. Each point corresponds to a particular gene/cell combination. The percent numbers above the boxplots give the total error in the proportion of the molecules counted in the dataset. The genes for which all of the methods gave correct results were excluded from the comparison.

**(B)** Dependency of the molecular abundance estimation errors on the number of molecules per gene. As in the Figure 2C of the main manuscript, the plots show the error in molecular count estimates in trimmed UMI data (expressed as a percentage of the molecular count), relative to the untrimmed *cluster-corrected* estimates (see Methods). Each point shows the median error (y axis) for all of the genes with the same untrimmed number of molecules (x-axis). Compared to Bayesian correction method, most methods underestimate the counts for highly expressed genes, even when trimmed by 1nt (9nt UMIs).

**Figure S6. UMI collisions on trimmed data.**

The plots show effect of different UMI collision adjustment approaches on the trimmed UMI data. The true number of molecules (estimated on untrimmed UMIs with *cluster* correction) is shown on the x axis, with the y axis giving the percent error for different lengths of trimmed UMIs (the error percentage is calculated relative to the untrimmed estimates). The lines were obtained using spline smoothing, as in Figure 2 of the main manuscript.

**Figure S7. Correlation of gene expression profiles within clusters.**

Similar to Figure 2D of the main manuscripts, the plots show distribution of within-cluster Spearman correlation of the estimated cell expression profiles for (A) mouse BMCs dataset, and (B) mouse pancreatic cells dataset. In the both cases, most of the correction methods result in similar improvements to within-cluster correlation.

**Figure S8. Initial labeling of high-quality cells based on cell size distributions.**

Two heuristic ways of assigning initial cell labels to high-quality and low-quality cells are shown.

**(A)** Shows scaled histogram, where the cells were binned based on the total number of molecules. For each bin, the y axis gives the number of molecules multiplied by the number of cells, so that the overall plot shows the distribution of molecular counts as a function of cell size. Such distributions tend to be bi-modal and the local minimum can be used to estimate the separation between the well-measured cell population (right) and low-quality/empty droplets (left). However, such pattern is not apparent in some datasets.

**(B)** Similar pattern can be obtained by changing the x axis to map molecular counts on the cell rank (with the largest cell being assigned the lowest rank).

**(C)** An alternative approach is to identify the size threshold based on the second derivative of the cumulative size distribution plot (see Methods). Here the y axis gives the total size of the cell, and x axis orders the cells by rank. The cells within the 25% of the identified thresholds (by rank) are initially labeled as Unknown (see Methods).

**Figure S9. Robustness of different classifiers to training errors.**

**(A)** As classification of low-quality cells depends on the ability to tolerate high level of errors in the initial training label assignment, we examined the robustness of different types of algorithms to artificially-added errors. Specifically, each classifier was first trained on the initial data, and the resulting labels were used for subsequent perturbations (to avoid simulation bias). Then 80% of the data was used to re-train the classifier, but a fixed percentage of class labels (x-axis) were swapped. The plot shows false positive and false negative rates (y axis) of different classifiers on the remaining 20% of the data.

**(B)** Analogous plot shows performance of the classifiers on the labels that were swapped in the 80% of the data that was used for training.

**(C)** Robustness of the classifiers to the initial labeling thresholds. The cell size borders used to assign the initial labels were widened by a fixed percentage (x axis), and the performance of the classifiers was compared using the labels, which the classifiers were able to predict based on the original borders.

**Figure S10. Difference between cell quality score and cell size rank.**

The plot illustrates the extent of disagreement between the labeling of large cells based on the cell size threshold (x axis) and the labeling based on the cell quality score. The measures were calculated by taking the rank-based labeling to be true. The optimal agreement is achieved at the minimum of the green curve. Mouse pancreas dataset was used.

**Figure S11. Comparison of the initial label assignments with the cell quality score predicted by the algorithm.**

A tSNE visualization shows all mouse pancreatic cells colored according to: (A) their initial (size-based) quality labels, and by (B) quality score returned by the KDE classification algorithm. Overall, the algorithm recognizes lower-size high-quality cells that are found within the major population clusters. Conversely, almost all cells outside of the major clusters are recognized with low (≤0.5) quality scores.

**Figure S12. tSNE visualization of cells selected based on the total number of molecules.**

Mouse pancreas cells are shown on t-SNE plots, colored by the published classification. Gray colors are used to show not clustered cells. Distribution of cell sizes identified cell size threshold separating 1524 top cells.

**(A)** Shows embedding of the top 75% of these cells (top 1197) which are the cells that are initial label of ‘good’ cells.

**(B)** Shows embedding of all 1524 cells.

**(C)** Shows embedding of all cells detected in the dataset.

All plots, including the most stringent size-selected (A) contain a grey cluster of unclassified cells. This artefactual cluster is removed when considering only cells with high quality score (>0.9, Figure 4B of the main manuscript).

